# Quantification of alterations in diffusion measures of white matter integrity associated with healthy aging

**DOI:** 10.1101/540443

**Authors:** Ciara J. Molloy, Sinead Nugent, Arun L.W. Bokde

## Abstract

This study aimed to evaluate the linear association of age with diffusion tensor imaging (DTI) measures of white matter such as fractional anisotropy (FA), mean diffusivity (MD), axial diffusivity (AD) and radial diffusivity (RD). We assessed patterns of overlap between linear correlations of age with FA with RD, MD and AD to characterize the process of white matter degeneration observed with ageing. 79 healthy adults aged between 18 and 75 took part in the study. The DTI data were based on 61 directions acquired with a b-value of 2000. There was a statistically significant negative linear correlation of age with FA and AD and a positive linear correlation with RD and MD, and AD. The forceps minor tract showed largest percentage of voxels with an association of age with FA, RD and AD, and the anterior thalamic radiation with MD. We found 5 main patterns of overlap: FA alone (15.95%); FA and RD (31.90%); FA and AD (12.99%); FA, RD and AD (27.37%); FA RD, and MD (6.94%). Patterns of overlap between diffusion measures may reflect underlying biological changes with healthy ageing such as loss of myelination, axonal damage, as well as mild microstructural and chronic white matter impairments. This study may provide information about causes of degeneration in specific regions of the brain, and how this may affect healthy brain functioning in older adults.

## Introduction

Diffusion tensor imaging (DTI) is a neuroimaging technique, which allows for non-invasive, in vivo, investigation of white matter [1–3]. DTI measures are based on random motion of water molecules, where within the brain diffusion of water is less restricted, or more isotropic, in areas of grey matter and CSF, and more restricted, or more anisotropic, in areas of white matter. When white matter structural architecture deteriorates, water molecules within white matter tissue become more isotropic, making DTI a useful tool for assessing atrophy [4–9].

The diffusion tensor is a 3×3 covariance matrix used to model diffusion within a voxel, in which there are 3 positive eigenvalues (λ1, λ2, λ3) and 3 orthogonal eigenvectors (ɛ1, ɛ2, ɛ3). The eigenvalues of the tensor give the diffusivity in the direction of each eigenvector. Together they describe diffusion probability using an ellipsoid, where the axes of the ellipsoid are aligned with the eigenvectors, and the major eigenvector (λ1) represents the principal diffusion direction.

Fractional anisotropy (FA), mean diffusivity (MD), radial diffusivity (RD) and axial diffusivity (AD) are the four main diffusion-based measurements of white matter structural architecture. FA measures the amount of diffusion asymmetry within a voxel, where a value of 0 is isotropic and is represented by a spherical ellipsoid with equal eigenvalues, and a value of 1 is anisotropic and is represented by an elongated ellipsoid with unequal eigenvalues. FA has been associated with the microstructural integrity of white matter [10, 11]. RD is a measure of perpendicular diffusivity and is an average of the two smaller eigenvalues (λ2+λ3/2). RD is thought to reflect myelination of white matter tracts, where an increase in RD would suggest loss of myelination [12–14], however white matter properties such as axonal diameter and density changes also affect RD [10]. AD represents parallel diffusivity and is a measure of the largest eigenvalue (λ1). AD has been linked to axonal integrity, where a decrease in AD would suggest axonal damage [15, 16]. AD decreases observed with axonal damage are thought to be caused by increases in debris from disrupted membrane barriers [17], whereas increases in AD have been suggested to reflect brain maturation [18]. However, increases in AD have also been linked to severe white matter injury when observed with changes in FA and RD [19, 20]. MD, also known as the apparent diffusion coefficient, is an average of all 3 eigenvalues in the tensor (λ1+λ2+λ3/3), and is therefore not independent of FA, RD and AD measures [10, 21]. It is thought to be a measure of membrane density. Together all four of these diffusion measures can distinguish distinct information about white matter microstructure.

The aging process coincides with changes in brain structure, brain function, and aspects of cognition such that they all decline as we get older. Many DTI studies have examined white matter structural changes that occur with aging [22–25]. Importantly, DTI studies of aging are consistent with post mortem findings of axonal and myelin damage observed in older adults, proving DTI to be a valuable in vivo measurement of white matter structural integrity [26, 27]. FA is the most commonly studied diffusion measure, and extensive decreases in FA with normal healthy aging have been reported in white matter across the brain including the body and genu of the corpus callosum (CC), the uncinate fasciculus (UF), the internal and external capsule, cingulum, fornix, the superior longitudinal fasciculus (SLF) and the inferior longitudinal fasciculus (ILF) suggesting widespread alterations in white matter occur with aging [20, 28–32]. Along with FA, age-related increases in MD have been reported in the genu and splenium of the CC [28], as well as the fornix, UF [31], IFOF, SLF, and ILF [32]. RD increases have been reported in the cingulum, CC, UF and ILF in older adults compared to younger adults [20, 21, 23, 33, 34]. Age-related changes in AD are less consistent, with findings of decreases in AD in the fornix, internal capsule, midbrain and cerebellum, but also reports of increases in AD in the CC [20, 23, 33]. However, other studies have also reported no changes in AD in older adults compared to younger adults [34–36], suggesting that perhaps AD changes are more subtle than RD. Further, studies showing greater RD changes compared to AD with aging indicate that myelination loss may be the main cause of degeneration of white matter with healthy aging [20, 25, 33, 34]. Longitudinal studies of older adults have shown global FA and MD changes, as well as FA, RD and AD changes in the genu of the CC within two years, suggesting that alterations in white matter can indeed occur over short periods of time with increased age [23, 37]. It has also been shown that white matter continues to mature in adulthood, with FA continuing to increase and MD continuing to decrease until between the ages of 20-42 years and 18-41 years respectively [38], stressing the importance of considering age-related changes across adulthood. Importantly maturation of white matter occurs at different rates across the brain, suggesting that perhaps the rate degeneration occurs differently in different tracts [23, 38].

Patterns of white matter changes within the brain have been examined to better understand and define the aging process, however conflicting findings have been reported. One pattern is the anterior-posterior gradient of degeneration, in which greater differences in FA in frontal regions compared to posterior regions of the brain; however although studies have reported this pattern, others have found no evidence of this [23, 33, 39]. Further, studies have even reported the opposite gradient pattern of larger FA decreases in posterior regions compared to frontal regions [33, 40]. A second proposed pattern is the superior–inferior gradient of white matter degeneration which postulates that superior white matter is more vulnerable to age-related changes [31, 41]. A study by Sullivan and colleagues (2010) reported both the anterior-posterior gradient and superior-inferior gradient of age-related changes in the brain indicating that perhaps age-related white matter changes may be more complex than one gradient of degeneration. A third pattern, described as the myelodegeneration hypothesis, proposes that myelin degeneration during aging occurs in the opposite direction of development [12–14]. This pattern would therefore be consistent with age-related changes in RD in white matter fibers that are last to myelinate, such as frontal and temporal associations fibers. Davis and colleagues (2009) reported age-related RD changes in the CC, cingulum and UF, along with larger age-related RD changes compared to AD, supporting the myelodegeneration hypothesis (Davis et al. 2009). However, again another study found no evidence of this pattern of age-related degeneration [42]. The inconsistent findings across the literature further suggest that these interpretations of white matter changes may be a simplification of the aging process, highlighting the need for a more thorough understanding of age-related effects on white matter.

While studies assessing age-related changes in diffusion measures separately have been informative, a combined analyses of FA, MD, RD and AD may provide a more comprehensive understanding of the underlying biological profile of the breakdown of white matter observed with aging [11, 24]. Indeed, a few studies quantifying specific patterns of changes in diffusion measures of white matter have proposed a more complex and region-specific interpretation of disruption observed with aging [20, 33]. Burzynska and colleagues (2010) quantified the frequency of overlap in age-related differences in measures of FA, RD and AD along with MD. A comparison of young and older adults established five main patterns of differences in diffusion measures observed with aging: (1) decreased FA with increased RD thought to reflect demyelination; (2) decreased FA with increased RD and MD thought to reflect chronic stages of white matter fiber degeneration; (3) decreased FA only thought to reflect mild microstructural alterations; (4) decreased FA with decreased AD thought to reflect acute axonal damage; and (5) decreased FA and AD with increased RD thought to reflect secondary Wallerian degeneration resulting in axonal loss and increased extracellular matrix tissue structures (Burzynska et al. 2010). Another study comparing younger and older adults assessed the overlap between FA, RD and AD measures and found three main patterns of age-related changes in diffusion measures; (1) decreased FA with increased RD only; (2) decreased FA and AD with increased RD; and (3) decreased FA, with both increased AD and RD (Bennett et al. 2010). Another study by Zhang and colleagues (2010) of young, middle-aged and older adults also looked at the concordance and discordance of FA age-related changes with MD, and RD with AD separately [43]. Unlike the two previous studies mentioned, this study did not focus the analysis on voxels with showing significant age-related FA changes. They found (1) regions of decreased FA with increased MD increases and related it to demyelination and axonal loss, (2) regions of MD increases with no FA decreases and interpreted it as inconclusive, (3) one region with FA decreases and no MD changes and related it to Wallerian degeneration, (4) a greater change in RD than AD and suggested it reflected demyelination and Wallerian degeneration, and finally (5) a one region with greater increase in AD than RD and reported this as inconclusive. These studies emphasize the importance of considering patterns of changes in diffusion measures to elucidate and define the specific profiles of age-related alterations in white matter regions across the brain.

The first goal of this study was to examine patterns of overlap between linear correlations of age with FA, RD, AD and MD in a cohort of young, middle-aged and older adults. Specifically, we quantified the frequency of patterns to establish the most prominent to least prominent pattern. Many previous studies assessing patterns of overlap between multiple diffusion measures have compared young and old adults [20, 33]. These studies have typically employed voxelwise statistical analyses of DTI data with a b-value of 1000; in the current study we used a higher b-value of 2000. We hypothesized that specific patterns of overlap between linear correlations of age with FA, RD, AD and MD would be observed, and that these different patterns may reflect specific regions of minor microstructural alterations, axonal damage, loss of myelination, and regions of more chronic white matter degeneration which are all thought to occur in aging. Examination of the patterns of overlap between linear changes in diffusion measures with age will complement previous studies examining differences between young and old.

## Materials and methods

### Participants

Seventy-nine, right-handed participants aged between 18 and 75 years (mean = 43.9 ± 18.33 SD) (male = 39) took part in the study. Exclusion criteria included permanent metallic objects in the body, a self-reported history of psychiatric or neurologic disorders or a history of a serious head injury. For participants over the age of 55 the Mini Mental State Exam (MMSE) was performed and participants with scores below 1.5 SD of the standardized scores were excluded from the study (Table 1).

**Table 1.** Participant demographics.

|  | Overall<br>(n=79) | Young<br>(n=33) | Middle<br>(n=16) | Older<br>(n=30) |
| --- | --- | --- | --- | --- |
| Age | 43.91(18.33) | 24.66 (5.32) | 46.25 (5.81) | 63.83 (4.46) |
| Gender (M:F) | 39:39 | 18:15 | 7:9 | 14:16 |
| NART | 33.04 (5.66) | 32.97(5.75) | 31.44 (5.29) | 35.40 (6.03) |
|  | 113.59 |  | 111.94 |  |
| NART IQ | (4.78) | 112.63 (3.92) | (4.84) | 115.53 (5.09) |
| MMSE | - | - | - | 29.33 (0.76) |
| WMH load (%) | 0.06 (0.10) | 0.04 (0.08) | 0.09 (0.15) | 0.06 (0.08) |

Measures are displayed as mean (±SD).

This study was approved by the Faculty of Health Sciences review board at Trinity College Dublin. Written, informed consent was obtained from all participants in accordance with the tenants of the Declaration of Helsinki. Participants were recruited through advertisements in local newspapers, church newsletters, and poster campaigns throughout universities in Dublin.

### MRI data acquisition

Whole-brain high angular resolution diffusion imaging (HARDI) data were acquired on a Philips Intera Achieva 3T MRI system (Best, The Netherlands) equipped with a 32-channel head coil at Trinity College of Institute of Neuroscience, Dublin. DWI data were collected using a single-shot echo-planar imaging sequence with echo time = 81 ms, repetition time = 14,556 ms, FOV 224mm, matrix 112×112, isotropic voxel of 2×2×2 mm, 65 slices of 2 mm thickness with no gap between slices. Diffusion gradients were applied in 61 directions, with b = 2000 s/mm2, and four b = 0 s/mm2 were acquired also. A 3D high resolution T1-weighted anatomical image was collected using a T1FFE (fast field echo) sequence with echo time = 3.8 ms, repetition time = 8.4 ms, FOV 230mm, 0.9×0.9×0.9 mm voxel size, 180 slices, for correction of EPI-induced geometrical distortions in the data. A T2-weighted fluid-attenuated inversion recovery (FLAIR) image was collected with echo time = 120 ms, repetition time = 11000 ms, 0.45×0.45 mm^2^ in-plane resolution, slice thickness = 5 mm, FOV = 230 mm.

### Estimated total intracranial volume

FreeSurfer image analysis suite version 5.3 (http://surfer.nmr.mgh.harvard.edu/) was used to perform automated volumetric segmentation of the T1-weighted anatomical image and extract estimated total intracranial volume (eTIV) for white matter hyperintensity load analysis [44].

### White matter hyperintensity volume

Automated segmentation of white matter hyperintensity (WMH) volume (ml) was calculated using the lesion segmentation toolbox (LST) version 2.0.15 (www.statistical-modelling.de/lst.html) for SPM12. T2-weighted FLAIR images were used for lesion segmentation, with T1-weighted anatomical images as a reference. Lesions were segmented using the lesion prediction algorithm, which consists of a binary classifier in the form of a logistic regression model trained on the data of 53 multiple sclerosis patients with severe lesion patterns obtained at the Department of Neurology, Technische Universität München, Munich, Germany. As covariates for this model a similar lesion belief map as for the lesion growth algorithm [45] this was used, as well as a spatial covariate that takes into account voxel specific changes in lesion probability. Parameters of this model fit are used to segment lesions in new images by providing an estimate for the lesion probability for each voxel. To describe WMH load quantitatively WMH volume was normalized by eTIV extracted from FreeSurfer analysis.

### Preprocessing of DWI data

DWI data were preprocessed using ExploreDTI (v4.8.4) software (www.exploredti.com/) (Leemans et al. 2009a). Data quality checks were performed, including visual inspection of correct orientation of gradient components and checking for gross artifacts. The data were corrected for motion correction, eddy current induced geometric distortions and for echo-planar imaging (EPI) deformations by co-registering and resampling to each subjects’ T1-weighted anatomical image [46]. Correction for each of these distortions was performed in one step. The B-matrix rotation was performed within this step to reorient the data appropriately [47].The default setting of robust extraction of kurtosis indices with linear estimation (REKINDLE) approach was utilized when the tensor model was applied to the data [48].

### TBSS analysis

FSL software (FMRIB Software Library-http://www.fmrib.ox.ac.uk/fsl/) [49] was employed to perform whole brain voxel-wise analysis of white matter using Tract Based Spatial Statistics (TBSS) [50]. FA diffusion images were extracted from diffusion data preprocessed using ExploreDTI.

A study specific template was created using all 79 subjects by aligning each subject’s FA image to every other subject’s FA image to identify the most representative subject. The study specific template chosen was then affine aligned to MNI standard space (included in FSL software package). Every other subject’s FA image was then warped into MNI space by combining the non-linear transformation to the chosen template FA image with the affine transformation to MNI space. This is done in one step to avoid resampling of the images twice. FA images from all subjects were then averaged to create the mean FA template and thinned to create a mean FA skeleton which represented the ‘skeletons’ of all tracts common to the group. We also investigated other diffusion measures extracted from data preprocessed in ExploreDTI such as MD, RD and AD. To do this we applied the same non-linear transformation and skeleton projection used for the FA images to the MD, RD and AD diffusion measures.

### Statistical Analysis

Voxel-wise statistics were performed using the randomise permutation-based inference tool for nonparametric statistical thresholding within FSL [51]. Permutation methods were employed as they are employed when the null distribution is not known. A linear correlation of age with each diffusion measure was performed, with WMH load, the National Adult Reading Test (NART) and gender included as covariates of no interest. The NART provides an estimate measure of premorbid intelligence [52]. As intelligence and gender differences in diffusion measures of white matter have been reported, we controlled for the effects of these measures in our analysis [32, 53]. It has also been suggested that WMH should be adjusted for when examining differences in healthy white matter [54]. In our supplementary material we provide the same analysis without any covariates included to compliment previous studies (see supplementary Table 1). The mean FA skeleton for the group was used as mask (thresholded to a value of 0.2) and the number of iterations was set to 10,000. The results were thresholded to a p value of <0.05 corrected for multiple comparisons across voxels using the threshold-free cluster enhancement (TFCE) option from the randomize permutation testing tool in FSL [55].

### Quantification of age-related white matter results

To quantify the size of the clusters of significant age-related linear correlations with diffusion measures across the whole brain, the percentage of voxels associated with aging was calculated by dividing the number of voxels showing statistically significant linear correlations with age for FA, MD RD and AD by the total number of voxels in the mean FA skeleton.

To quantify the age-related linear correlations with diffusion measures within white matter tracts ROIs were created using the JHU Tractography atlas. Specifically, each ROI was masked, thresholded to above 10, and multiplied by the mean_FA_skeleton to create masks of total voxels within each tract. Statistically significant results for each diffusion measure were multiplied by tract ROIs to create masks of voxels showing linear correlations with age. The percentage of age-related voxels within tract ROIs was calculated by dividing FA, RD, AD and MD masks of linear correlations with age by the total tract masks.

### Quantification of overlap between diffusion measures

The overlap between voxels showing age-related linear correlations with FA, MD, RD and AD was assessed. First the TBSS results showing significant linear correlations of age with FA, MD, RD and AD were masked. Then the FA mask was multiplied by each of the MD, RD and AD masks to create masks of the overlap between diffusion measures. The percentage of overlapping voxels was calculated by dividing each of the overlap masks by the FA mask.

## Results

All statistically significant linear correlations of diffusion measures with age reported in the results include WMH load, NART scores and gender as covariates to account for any changes in diffusion measures of white matter that may be related to them.

### Age-related linear correlations with diffusion measures

We quantified the age-related linear correlations in each of the diffusion metrics by calculating the percentage of voxels showing statistically significant linear correlations with age, with NART and gender included as covariates (Table 2). Age-related statistically significant negative linear correlations with FA were observed in 49.69% of white matter voxels, statistically significant positive linear correlations with RD were observed in 40.00%, statistically significant positive linear correlations with MD in 7.73%, statistically significant negative linear correlations with MD were observed in 0.41%, statistically significant positive linear correlations with AD in 0.31%, and statistically significant negative linear correlations with AD in 31.53% of white matter voxels. There were no statistically significant positive linear correlations of FA with age or statistically negative linear correlations of RD with age.

**Table 2.**
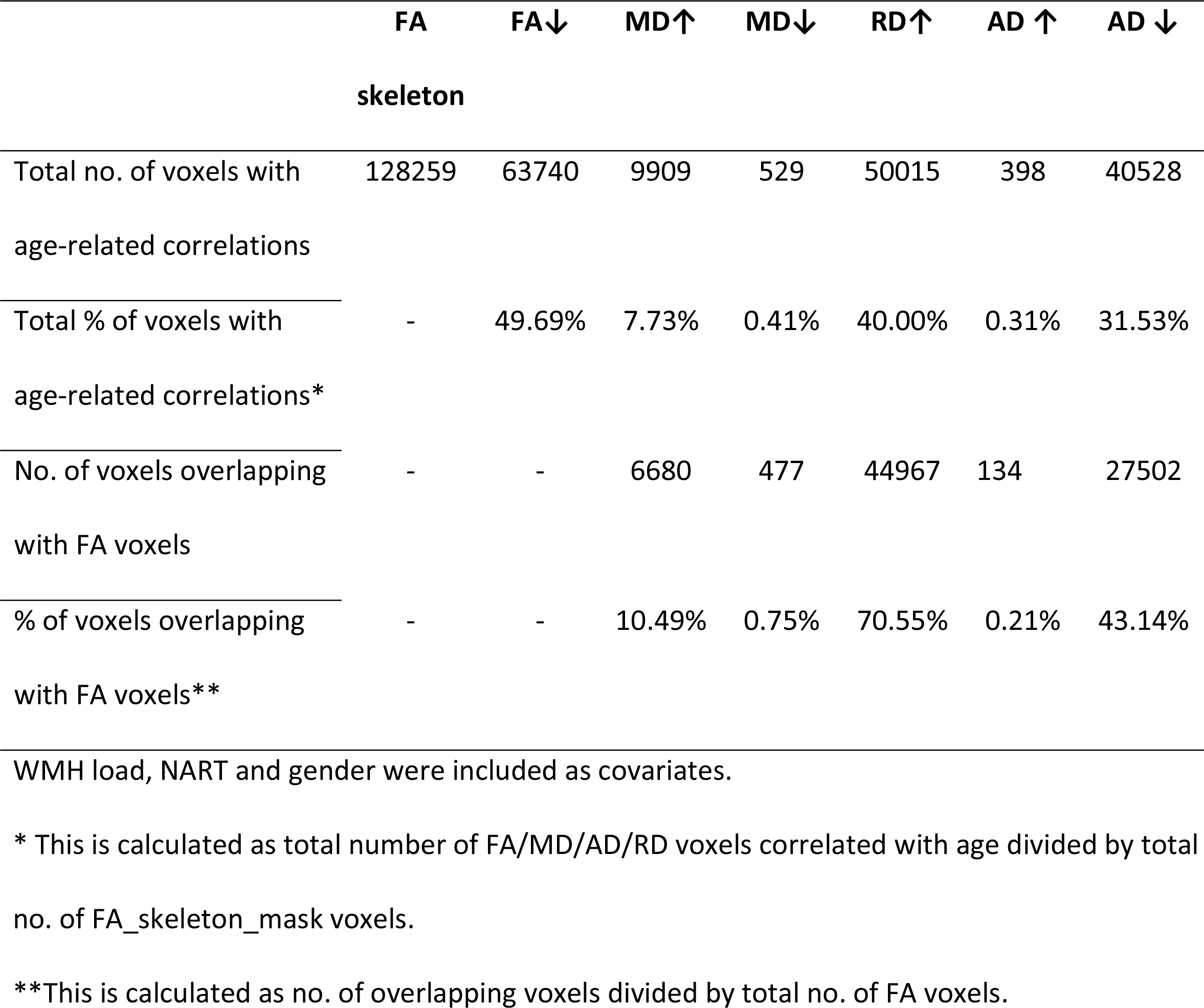
Total number and percentage of voxels correlated with age, and the overlap of MD, RD and AD with age-related FA voxels.

We then assessed percentage of white matter regions affected by aging using the JHU white matter tractography atlas (Table 3). FA, RD and AD all showed the greatest percentage of voxels correlated with age in the forceps minor (90.28%, 81.73% and 71.17% respectively), whereas MD showed the greatest percentage of voxels correlated with age in the right anterior thalamic radiation (16.22%). All tracts showed some percentage of voxels with age-related linear correlations with FA and RD; however, the cingulum hippocampus showed the smallest percentage (FA: 2.71% and 2.33% for left and right respectively; RD: 0.78% and 3.10% for left and right respectively). For MD and AD, there were no voxels with age-related statistically significant positive or negative linear correlations in the cingulum hippocampus.

**Table 3.**
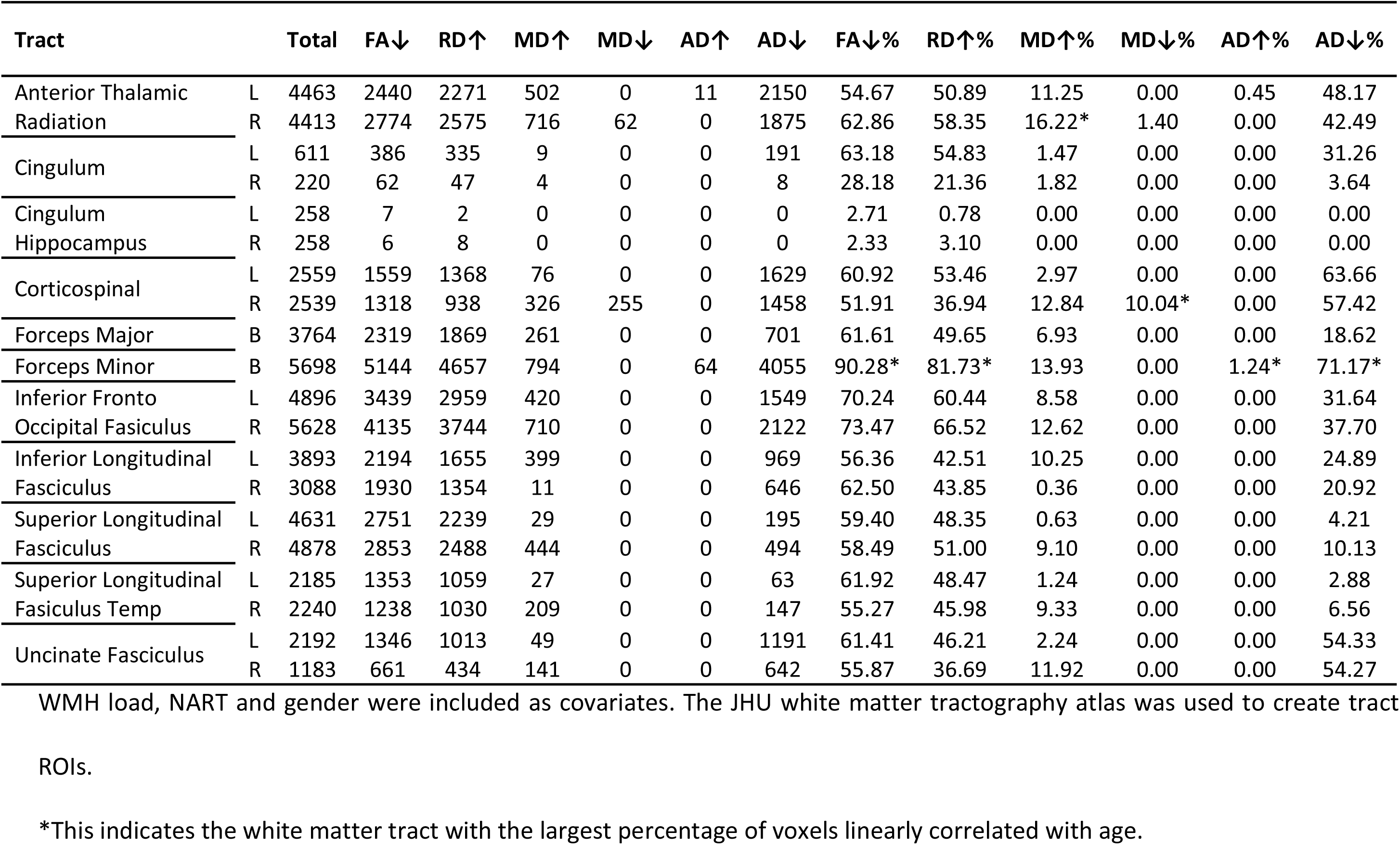
Number and percentage of voxels showing age-related correlations with diffusion measures.

### Overlap between age-related linear correlations with diffusion measures

Within FA voxels showing significant negative age-related linear correlations, we calculated the percentage of voxels also showing statistically significant linear correlations of age with each of the diffusion measures of MD, RD and AD (Fig 1 and Table 4). Voxels showing statistically significant positive age-related linear correlations with RD overlapped with 70.55% of voxels showing statistically significant negative age-related linear correlations with FA. Positive and negative statistically significant linear correlations with AD overlapped with 0.21% and 43.14% of voxels showing statistically significant negative age-related linear correlation FA respectfully. Positive and negative statistically significant linear correlations with MD overlapped with 10.49% and 0.75% of voxels showing negative age-related statistically significant linear correlations with FA respectfully.

**Table 4.**
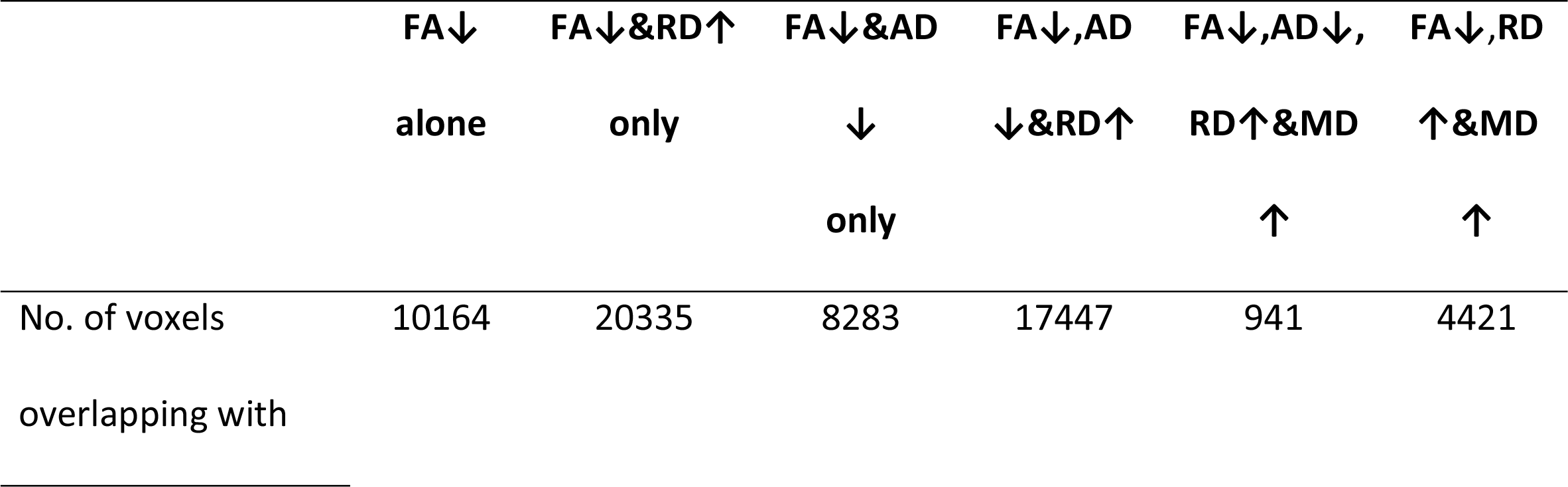
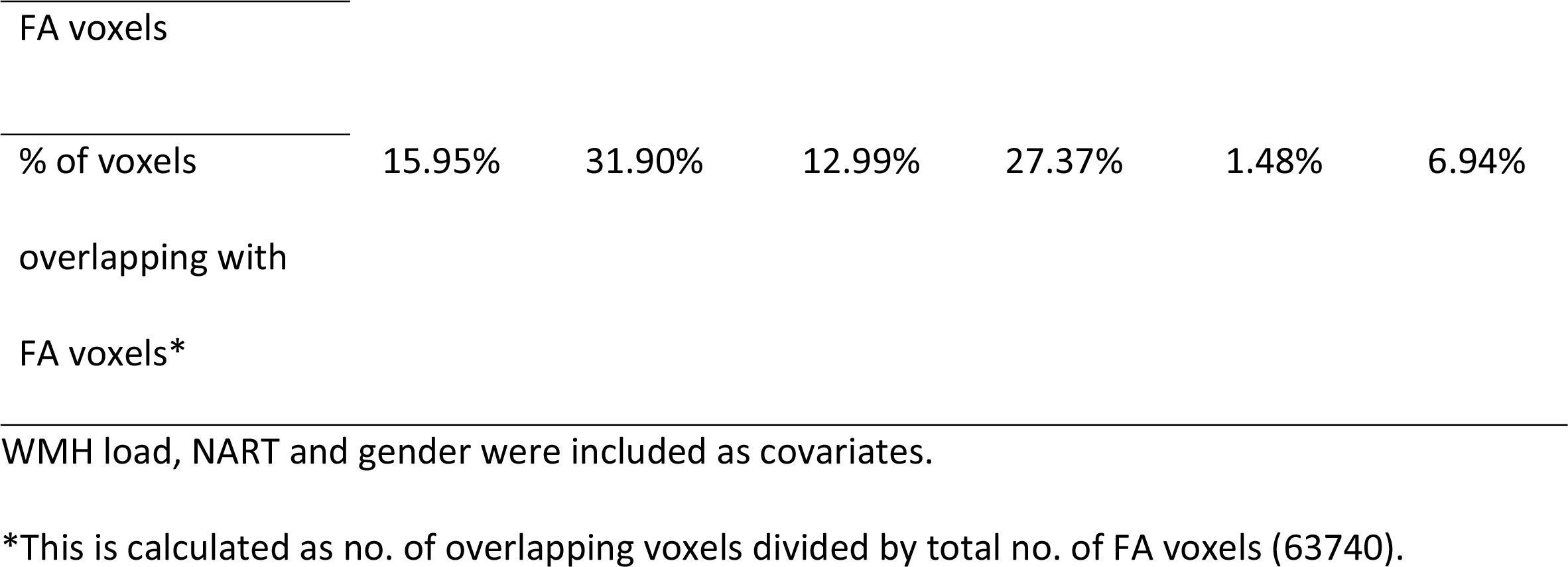
Total number and percentage of voxels correlated with age showing specific patterns of overlap between diffusion measures.

**Fig 1.**
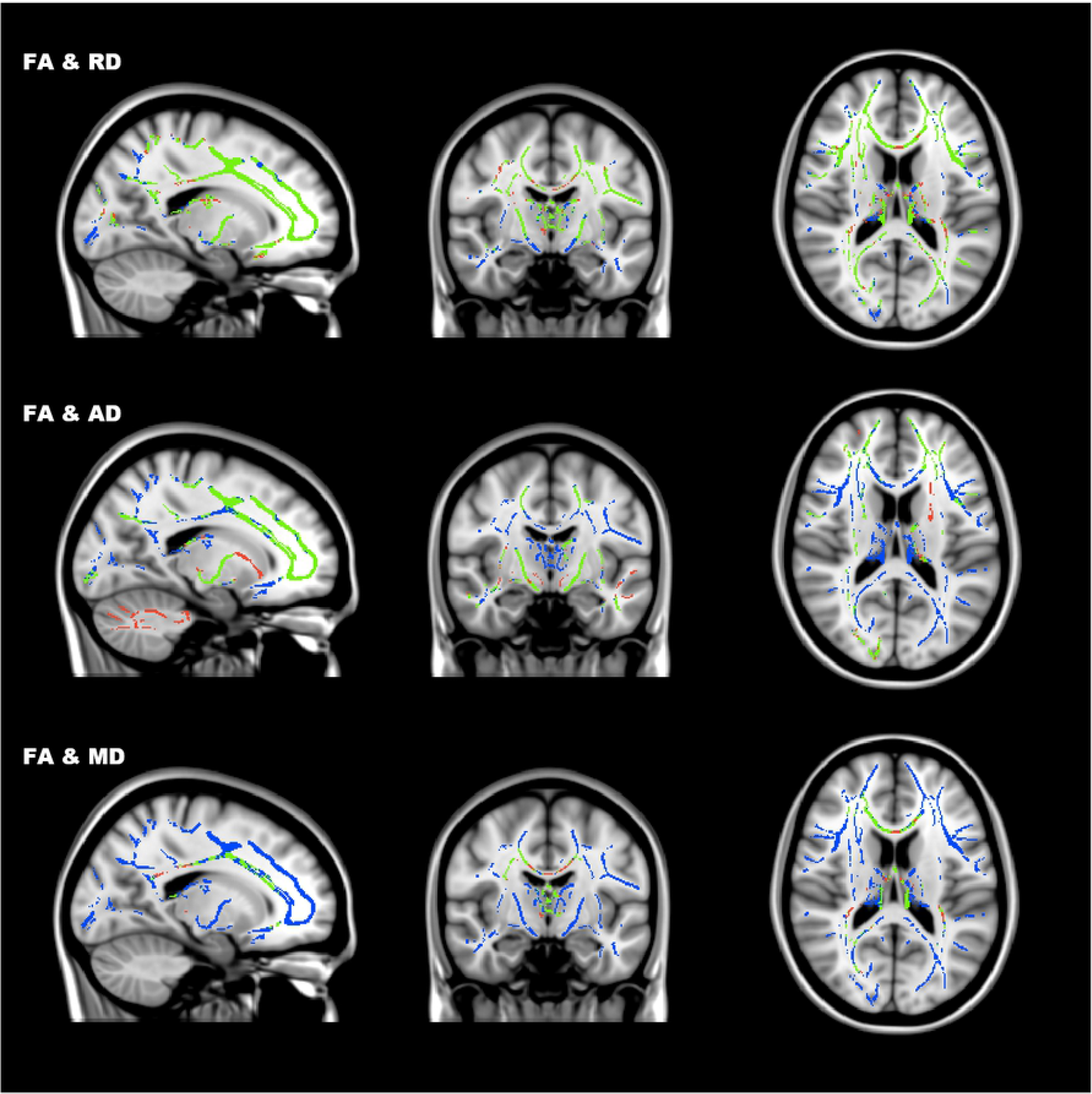
Overlap between statistically significant linear correlations of age with FA and RD, AD and MD. Age-related positive correlations with RD overlapped with 69% of age-related negative correlations with FA; AD positive correlations overlapped with 43% of FA correlations; and MD with 8%. Images are displayed in radiological convention (i.e. left=right). FA = blue; RD, AD, MD = red; Overlap = green.

We further identified 5 main patterns of overlap between diffusion measures in white matter regions showing age-related negative linear correlations with FA, and a sixth pattern in very few voxels (Fig 2 and Table 4):

**Fig 2.**
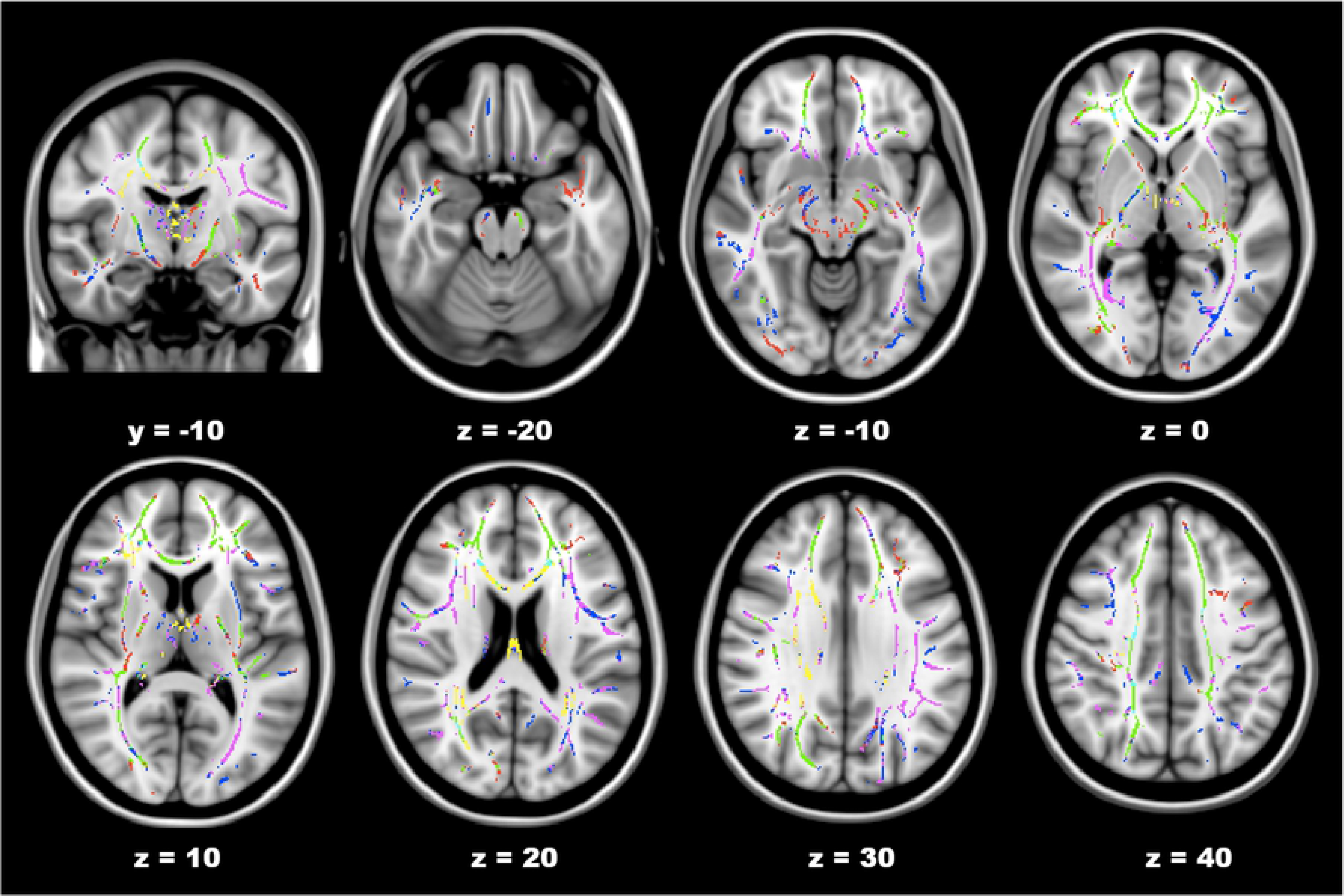
Patterns of overlap between diffusion measures within voxels showing age-related correlations with FA. Images are displayed in radiological convention (i.e. left=right). FA alone = blue; FA & RD = pink; FA & AD = red; FA, RD&AD = green; FA, RD & MD = yellow; FA, RD, AD & MD = light blue.

1. FA and RD only pattern: statistically significant positive age-related linear correlations with RD alone showed the greatest percentage of overlapping voxels with FA across the brain (31.90%). This was the main pattern observed in the forceps major, right and left SLF, right and left ILF, right and left IFOF and right cingulum. This pattern was also observed in the right and left anterior thalamic radiation, and smaller clusters of the right and left UF, corticospinal tract, and forceps minor and body of the CC, and right anterior limb of the internal capsule.

2. FA, RD and AD pattern: statistically significant negative age-related linear correlations with AD and statistically significant positive age-related linear correlations with RD showed the second greatest percentage of overlapping voxels with FA linear correlations (27.37%). This was the main pattern observed in the forceps minor, right and left UF, right and left corticospinal tract, left posterior limb of the internal capsule, and left cres of the fornix. This pattern was also observed in a large number of voxels in the left and right IFOF, the left retrolenticular part of the internal capsule, and smaller number of voxels in the right and left ILF, SLF, and the forceps major.

3. FA only pattern: statistically significant negative age-related linear correlations with FA alone showed the third greatest percentage of overlapping voxels (15.95%). This was not the main pattern in any tract ROI. It was observed in the right and left SLF and temporal SLF, as well as small regions of the right and left anterior thalamic radiation, cingulum, forceps major and minor (body and left splenium of the CC), right and left IFOF, ILF and UF, right internal capsule and the left external capsule.

4. FA and AD pattern: The fourth pattern observed was age-related statistically significant negative linear correlations with AD and FA (12.99%). This was the main pattern observed in the right UF along with the FA, AD and RD pattern. It was also observed in clusters of the forceps major and minor, right and left anterior thalamic radiation, right and left corticospinal tract, right and left IFOF, right and left ILF as well as the right SLF and left UF, small clusters of the of the right external capsule, and the retrolenticular part of the right internal capsule, as well as right and left internal capsule.

5. FA, RD and MD pattern: The fifth pattern observed was age-related statistically significant positive linear correlations with RD and MD, with statistically significant negative linear correlations with FA (6.94%). This pattern was the main pattern in the fornix (column and body). It was also found in small regions of the forceps minor (also genu and body of the CC), and the right and left anterior thalamic radiation, right anterior IFOF and posterior IFOF/ILF, right corticospinal tract, forceps major, right and left SLF, and right and left anterior thalamic radiation.

6. FA, RD, AD, and MD pattern: The sixth pattern observed was age-related statistically significant negative linear correlation with FA and AD, with statistically significant positive linear correlations with MD and RD 1.48%). This pattern was found in a small cluster of voxels in the right IFOF, right and left UF, the forceps minor and body of CC, and right SLF.

## Discussion

### General findings

This study aimed to quantify the linear effect of age on FA, RD, AD and MD diffusion measures of white matter integrity in a cohort of young, middle aged and older healthy adults, after adjusting for white matter hyperintensity (WMH) load, premorbid IQ (NART) and gender. Our findings showed negative linear correlations of age with FA and AD, as well as MD in a small number of voxels. We also found a positive linear correlation of age with MD and RD, as well as AD in a small number of voxels.

The findings of negative correlations of FA with healthy aging in many regions of the brain is in accordance with decreased FA in older adults compared to younger adults in many previous studies [21, 23–25, 29, 33, 39, 56]. When we quantified the number of voxels linearly correlated with age our findings showed that FA was statistically significant correlated with age in approximately half of the brain’s white matter, higher than all other diffusion measures. This was similar to the findings from a study by Burzynska and colleagues (2010) showing age-related decreases in FA in 53% of voxels. We further showed that the tract with the highest percentage of voxels showing age-related linear correlations with FA was the forceps minor, which was also true for RD and AD, suggesting that this tract may be greatly affected by typical aging. Interestingly age-related associations with FA were found in approximately 50% of voxels or more in all tracts except the right cingulum, and the right and left cingulum hippocampus, which is consistent with the parahippocampal segment of the cingulum. This points to generally extensive global effects of healthy aging on white matter integrity. The small number of voxels showing associations of age with FA and RD in the hippocampal branch of the cingulum, and the lack of any association with AD or MD indicates this region of white matter is less affected than other white matter tracts. Similar to our findings, a study comparing young and older healthy adults found differences in the subgenual branch of the cingulum but no change in any diffusion measures in the parahippocampal segment of the cingulum [57].

Our analysis revealed a larger percentage of voxels showing linear correlations of age with RD (positive = 40.00%; negative = 0%) than with AD (positive = 0.31%; negative = 31.53%) and MD (positive = 7.73%; negative = 0.41%) indicating more widespread changes in RD across the brain with aging. In line with our initial findings, greater changes in RD compared to AD with aging has been reported previously suggesting that healthy aging is more affected by myelination loss than axonal damage across the brain [20, 33, 34, 43].

Unlike Burzynska and colleagues (2010) we found only a small number of voxels with negative linear correlations of age with MD. However, this is a less consistent finding in the literature, with more studies reporting an increase in MD with aging [21, 28, 31, 32]. We also found a small number of voxels showing a positive linear correlation of age with AD. Both increases and decreases of AD with age have been reported [20, 23, 33, 57]. Animal studies have attributed decreases in AD to acute axonal injury [13, 58]. Human diffusion studies have observed AD increases with traumatic brain injury, resulting in more chronic and irreversible white matter damage [59]. Taking this into account, perhaps the cohort in the current study had less chronic axonal damage than previous studies. However, another explanation for AD increases may be due to loss of fiber coherence in regions of crossing fibers, which is a common issue with the diffusion tensor model [60].

To further inform age-related changes in white matter integrity we next assessed patterns of overlap between diffusion measures significantly linearly correlated with age. Specifically, we examined patterns only within voxels showing significant negative linear correlations of age with FA. That RD overlapped with ~70% and AD with ~43% of voxels showing age-related linear correlations with FA, is indicative of a strong association of both RD and AD with aging in our cohort. The greater percentage of overlap between FA and RD than FA and AD, further suggests that loss of myelination may be a major contributor to degeneration observed with healthy aging [33, 34].

While our results showed multiple patterns of overlapping diffusion measures within tracts, a main pattern was observed for most tracts. Multiple patterns of overlap within tracts may reflect vulnerability in specific regions of the brain, rather than white matter tracts as a whole. It is also possible that examination of patterns of overlap reveals regions of white matter tracts showing different stages of white matter changes observed with aging, or simply that perhaps degeneration is not consistent across the brain, which would not be surprising given that some studies have suggested different gradients of changes in diffusion of white matter integrity with aging [31, 34, 61]. However, given that a main pattern was observed in many tracts including the forceps minor, forceps major, left and right SLF and ILF, our findings suggest there may be a specific profile of age-related changes in major white matter fiber bundles.

### The FA and RD pattern

The most prominent pattern observed was a linear correlation of age with FA and RD, but not AD, which has been thought to reflect mainly the process of myelination loss in white matter tracts [13, 14, 20, 33]. Histology investigations have indicated that white matter atrophy is a major causal factor of brain degeneration with aging, supporting DTI studies [27]. Further, loss of myelination in the CC, a highly myelinated white matter bundle, for which the genu and splenium sub regions are consistent with the forceps major and minor respectively, has been shown in older adults [27]. Consistent with this, in the current study the FA and RD pattern was largely observed in the forceps major, along with the IFOF, ILF and SLF. Similarly, Burzynska and colleagues’ (2010) study comparing older and younger adults also reported this pattern as the dominant pattern in forceps major [20].

Differences in white matter integrity in the SLF with aging have been reported previously [62], as well as the FA and RD pattern of age-related changes in the SLF [33]. Although the FA and RD pattern was the prominent pattern in the SLF, it should be noted that we also found voxels showing the FA alone pattern in both the left and right SLF as well as the FA, AD and RD pattern and FA, RD and MD pattern in a small number of voxels of the right SLF.

For the ILF and IFOF findings, it is important to note that they are known to overlap partly, so diffusion measures may not distinguish between these tracts very well along some regions [63]. We also found some clusters of voxels within both tract ROIs showing other patterns. Previous studies of age-related changes in the IFOF are mixed. One previous study examining frontal lobe white matter connections, did not find age-related linear correlations with diffusion measures in this tract [62]. However, differences in FA have been reported in older adults compared to young [31, 32, 64]. It is possible that these discrepancies across studies may reflect differences in DTI acquisition parameters or analysis techniques. Overall, it seems that loss of myelination may indeed explain a large proportion of white matter degeneration observed with aging.

### The FA, RD and AD pattern

The FA, RD and AD pattern has been linked to axonal damage, gliosis and subsequent increases in extracellular tissue structures [3, 20]. While decreases in AD have been reported in studies of ischemic stroke, which have been suggested to result in gliosis. This was the main pattern in many tracts with frontal lobe connections such as the forceps minor, anterior thalamic radiation, UF, and a large proportion of the IFOF, as well as other white matter fibers such as the corticospinal tract and the cres of the fornix, and cingulum.

The frontal lobes have been suggested to be sensitive to aging [20, 33, 41]. Although studies have indicated that frontal regions may be more vulnerable to age-related decline than posterior regions [41], a longitudinal study assessing rate of change in diffusion measures did not find evidence of the anterior-posterior gradient over a 2-year period [23]. Our findings of widespread age-related degeneration in white matter, with multiple patterns across the brain also indicate that the pattern of degeneration may be more complicated. For example, although the FA, RD and AD pattern was the main pattern in the UF, some voxels showed the FA and RD pattern, as well as the FA and AD pattern. Interestingly, a previous study assessed diffusion measures separately and found changes in FA, RD, AD and MD with aging in the UF, suggesting that an overlap in age-related changes could have also been observed in the UF [31].

The cingulum, which connects the frontal lobe with parietal and temporal regions such as the hippocampus and parahippocampus, showed the FA, RD and AD pattern in the right and left cingulum, as well as a similar number of voxels with the FA and RD pattern in the left cingulum [65–67]. However, only a very small number of voxels within the cingulum hippocampus ROI were associated with aging suggesting that this tract was not affected by age. Other studies have also observed differences in FA and RD only in parts of the cingulum, such as the superior portion but not posterior or inferior region [31], and the subgenual branch but not the parahippocampal segment of the cingulum [57].

This FA, RD and AD pattern was also reported in the crus of the fornix by a previous study comparing younger and older adults, suggesting this small white matter structure is vulnerable to healthy aging possibly due to axonal damage and gliosis [20]. Our findings suggest that axonal damage and increases in extracellular matrix tissues may be another major cause of brain white matter changes that occur with the normal healthy aging process.

### The FA alone pattern

The FA alone pattern has been suggested to reflect mild microstructural changes, such as fiber structure loss without any major tissue loss, where these decreases in FA in the absence of MD increases may be due to subtle changes in RD and AD [20, 43]. Although this pattern was observed throughout the brain, it was not the most prominent pattern in any white matter tract. This indicates that FA overlapped with other diffusion measures in a large proportion of voxels showing age-related linear correlations. Consistent with the straight gyrus reported by Burzynska and colleagues (2010), we also observed the FA alone pattern in the inferior frontal gyrus. Overall the current study findings suggest that these less severe microstructural changes occur across the brain with aging. The fact that FA showed the largest percentage of affected voxels, indicates that perhaps FA alone mild may changes occur first, may then be followed by changes and/or greater changes in other diffusion measures. Future studies comparing young and middles aged adults or longitudinal designs are needed to investigate this further.

### The FA and AD pattern

The FA and AD pattern was the fourth pattern we observed which has been linked to acute axonal damage, such as axon swelling and fragmentation, and therefore distinguishing it from the FA, RD and AD pattern [20]. Although this pattern was observed in a proportion of voxels in many tract ROIs, it was not the dominant pattern in any tract. This indicates that correlations of AD with age were mainly observed in conjunction with FA and RD, and not with FA alone. This would not be surprising given that most studies to date show greater RD changes than AD, and oThis pattern was previously reported by Burzynska and colleagues (2010), who also showed some areas of overlap with MD within this pattern. Interestingly, another study did not report this pattern, but instead only observed AD changes in conjunction with FA and RD, further suggesting it is not a common pattern associated with the aging brain [33].

### The FA, RD and MD pattern

The pattern of FA, RD and MD has been suggested to reflect chronic white matter fiber degeneration [20]. This pattern has been observed in While earlier studies have reported differences in all of three of these measures with aging, few have assessed their specific regional overlap [20, 31, 64]. Findings from our study indicated that chronic WM degeneration was observed in the CC, mainly in the regions of the body and genu as well as the body of the fornix. Diffusion imaging studies have consistently reported white matter structural changes in the CC with aging, particularly in the genu and body [20, 31, 33, 34, 62]. The CC is important for interhemispheric communication, therefore chronic white matter impairments in this white matter bundle could result in deficits in cognitive functions requiring cross communication between hemispheres [68]. For example, many functional MRI studies have shown bilateral activation in older adults compared to unilateral activation observed in younger adults, which has been suggested to be a compensatory mechanism for declining networks and additional brain recruitment for task performance [69–73]. It is plausible that this bilateral functional activation often reported within frontal lobes in older adults may result from white matter alterations in the CC affecting inter-hemispheric communication.

The fornix has previously been reported to show age-related structural changes [20, 31, 61, 74]. In contrast to our findings, a previous study found an overlap between FA, RD and MD along with increases in AD in the body of the fornix [20], and another study that only assessed FA, RD and AD, also found and overlap between all three measures in the fornix. It is important to note that the fornix is located close to the ventricles, so measures within this structure may be influenced by cerebrospinal fluid pulsation [74].

Although we found this pattern in some other white matter brain regions, again it was not the main pattern, suggesting that chronic white matter degeneration is not linearly changing across the brain with healthy aging.

### White matter hyperintensity load

A recent study by Svärd and Colleagues (2017) showed that WMH load can impact FA and MD measures in healthy elderly adults, and when comparing to prodromal Alzheimer’s disease, suggesting it may be essential to control for in any study assessing age effects on diffusion measures of white matter [54]. Given that WMH load increases are observed with aging, it is possible that some white matter differences between young and older healthy adults may be attributable to disparate WMH load [75, 76]. Therefore, it may be important to measure WMH load when assessing healthy aging, as well as neurodegenerative diseases to better understand white matter changes that are specific to “normal appearing white matter”, and not driven by white matter hyperintensities [54]. We performed the analysis in the same cohort without including WMH load as a covariate (i.e. only NART and gender covariates were included in correlation analysis) and found that adjusting for WMH load did not affect the results greatly in this study. For this analysis one extra subject was included (n = 80; mean age = 43.65 ± 18.36 SD), and the percentage of voxels correlated with age were minimally affected for each diffusion measure: FA (positive = 0%; negative = 50.17%), RD (positive = 39.01%; negative = 0%), MD (positive = 6.11%; negative = 0%) and AD (positive = 0.03%; negative = 31.63%) (see S1 Table). The main difference was that no negative correlations of age with MD were observed. The patterns of overlap, and percentages were also largely maintained (see S2 Table). It is worth noting that with the exception of one subject, all WMH loads were in the lower range for all subjects suggesting that our older adults were indeed healthy, and that perhaps subtler effects would be expected in the current cohort. Nevertheless, few studies have taken WMH load into account when assessing aging, and to our knowledge, none have assessed its effect on percentages of voxels linearly associated with age or patterns of overlap between diffusion measures. It is possible that discrepancies within aging studies may be accounted for, or at least partially explained by white matter hyperintensity load effects on DTI measures. Future studies of aging, or age-related diseases are needed to confirm the extent to which WMH load effects DTI measures in various tracts throughout the brain.

### DTI data acquisition insights

In the literature to date many TBSS studies of aging have used b-values of less than 1500, and the two main studies we compared to by Burzynska and colleagues (2010), and Bennett and colleagues (2010) both used b-values of 1000, whereas this study utilized diffusion weighted data with a b-value of 2000. Increasing the b-value can be advantageous; for example, it increases the brightness of white matter which may be beneficial to segmentation of gray and white matter [77]. An increase in b-value from 1000 to 3000 has also been shown to result in noisier images, loss of signal, and hyperintense white matter compared to gray matter [78]. The findings from this study of statistically significant negative linear correlations of age with FA are consistent with the literature, as well as the linear correlations with RD and AD. However, some dissimilar findings were found, mainly with MD. It has been shown that increases in the b-value results in decreases in the apparent diffusion coefficient but not FA measures, and changes in the diffusion properties of gray and white matter [77, 79]. Discrepancies between studies may indeed be explained by differences in data acquisition parameters. Further research focusing on differences in scan acquisition and analysis techniques is needed to test this specifically. Overall, this study found that a b-value of 2000 is still sensitive to detection of age-related changes in diffusion measures of white matter in the brain, using a widely used analysis method. We also found very similar patterns of overlap between FA, RD and AD diffusion parameters, observed in previous studies. Any discrepancies may be in part due to differences in acquisition parameters and/or various processing and analysis techniques.

### Limitations

DTI-based measures of axonal integrity and myelination should be interpreted with caution as these are indirect measures of white matter structural integrity. The TBSS technique also has limitations. It is known that 90% of the brain white matter consists of crossing fibers, however the diffusion tensor has difficulty in accounting for these crossing fibers [80]. Therefore, DTI measures may not be true representations of fiber integrity in regions of crossing fibers. Anatomical specificity inaccuracies of the TBSS analysis approach have also been reported, as well as image registration [81]. Despite these limitations, this analysis has been utilized by many studies. Importantly DTI studies of aging have reproduced results of post-mortem studies of the brain proving it to be a useful technique.

## Conclusion

This study quantified patterns of age-related linear correlations with diffusion measures of white matter in a cohort of young, middle-aged and older adults. Overall this study found that healthy aging has a profound impact on white matter. The patterns of age-related linear correlations with diffusion measures provides crucial information about possible underlying causes of degeneration of white matter, and how the various patterns of decline may be region specific. These findings may have important implications for connectivity between brain regions and the cognitive functions they support in healthy aging. The study emphasizes the importance of considering patterns of white matter changes in DTI measures across adulthood, while controlling for variables that may affect assessment of changes white matter with healthy aging. This may help to better understand specific biological profile contributing to white matter deterioration which may influence brain function in healthy aging.

## Acknowledgments

The authors would like to thank the participants who donated their time and energy to our research. The authors would also like to thank Sojo Josephs and the Trinity Centre for High Performance Computing (TCHPC) for their support throughout the project.

## Supporting Information

**S1Table. Total number and percentage of voxels correlated with age, and the overlap of MD, RD and AD with age-related FA voxels**. NART and gender were included as covariates.

**S2 Table. Total number and percentage of voxels correlated with age showing specific patterns of overlap between diffusion measure.** NART and gender were included as covariates.

